# Heart rate of the blue mussel Mytilus edulis in a simulated intertidal environment

**DOI:** 10.64898/2026.09.28.754745

**Authors:** Duy Nghia Pham, Inna M. Sokolova

**Affiliations:** Department of Marine Biology, Institute of Biological Sciences, Faculty of Mathematics and Natural Sciences, University of Rostock, Rostock, Germany; Department of Maritime Systems, Interdisciplinary Faculty, University of Rostock, Rostock, Germany

**Keywords:** Multiple stressors, Cardiac activity, Oxygen debt, Thermal response, Extreme event, Biomonitoring

## Abstract

Despite being a topic of early experimental studies, the effects of immersion-emersion cycles and temperature on the heart rate of intertidal *Mytilus* mussels have not been fully characterized. Using a recently released intertidal chamber paired with our own software package, we conducted a multiday experiment and continuously monitored the heart rate of the blue mussel *Mytilus edulis*. Under normal conditions, we detected a subtle spike in heart rate upon emersion before the eventual decline, as well as an overshoot upon reimmersion. Additionally, heart rate was sensitive to temperature changes during immersion but less responsive during emersion. These patterns were markedly altered following an unintended prolonged emersion event, which led to the loss of the emersion spike, an extended reimmersion overshoot, and heightened thermal sensitivity on top of an overall elevation in heart rate. However, the heart rate gradually returned to the pre-event baseline over the following days, demonstrating the significant resilience of the mussels. Our study highlights the potential of high-resolution physiological data in assessing the effects of environmental stressors on coastal ecosystems.

## 1 Introduction

The intertidal zone is the coastal area alternately submerged at high tide and exposed at low tide (Trujillo and Thurman, 2017). This repeated exposure to air subjects resident marine animals to multiple abiotic stressors, including desiccation and temperature fluctuations (Leeuwis and Gamperl, 2022). Despite these challenges, sessile species such as mussels of the genus *Mytilus* remain dominant in rocky intertidal communities across temperate and polar regions (Gaitán-Espitia et al., 2016; Gosling, 2021). How these mussels respond to their dynamic environment is a longstanding topic of interest in ecophysiology (Bayne, 1976). Today, this foundational knowledge is increasingly important for assessing the effects of intensifying environmental stressors, such as chemical contamination and extreme weather events (Achterberg et al., 2025).

Early experimental studies on intertidal *Mytilus* mussels have established an understanding of several key physiological parameters, particularly heart rate (Table 1; Schlieper, 1955). Generally, mussels responded to the transition from immersion to emersion by closing their valves, followed by a drastic decrease in heart rate that remained low throughout emersion. Upon reimmersion and valve reopening, the heart rate recovered and often temporarily exceeded the rate observed during the previous immersion, a phenomenon widely known as “reimmersion overshoot” (e.g., Coleman, 1972). However, much less documented is the brief increase in heart rate at the onset of emersion (Bayne et al., 1976; Zenone et al., 2009), herein termed “emersion spike”. Since the emersion spike has also been reported in other intertidal bivalves, including *Cerastoderma* cockles (Boyden, 1972) and *Crassostrea* oysters (de Carvalho et al., 2022, 2026), its presence in *Mytilus* mussels deserves further investigation.

**Table 1:** Chronological summary of experimental studies on the effects of transitions between immersion and emersion on the heart rate of *Mytilus* mussels. The emphasis is on the transition protocol (I: immersion and E: emersion), the detection of an emersion spike or reimmersion overshoot of heart rate (explicitly stated or visually apparent), and the temperature regime applied during emersion.

| Research and data source | Species and location | Transition protocol | Emersion duration | Emersion spike | Reimmersion overshoot | Temperature regime |
| --- | --- | --- | --- | --- | --- | --- |
| Schlieper (1955): figs. 4-6 | *M. edulis*, Sylt, Germany | I-E-I | 3 h | No | Yes | Constant |
| Helm and Trueman (1967): fig. 3a | *M. edulis*, Millport, Scotland | I-E-I | 3 h | No | Yes | Constant |
| Coleman and Trueman | *M. edulis*, Millport, | I-E-I | ~17 h | No | Yes | Constant |
| (1971): fig. 3a | Scotland |  |  |  |  |  |
| Coleman (1972): figs. 26, 28-32 | *M. edulis*, Conwy, Wales and Morecambe, England | I-E-I | 2-72 h | No | Yes | Constant |
| Coleman (1973): figs. 5b, 6 | *M. edulis*, Morecambe, England | I-E-I and I-E-I-E-I | 6 h, 16 h | No | Yes | Variable |
| Bayne et al. (1976): figs. 6-8 | *M. californianus*, San Juan Island, USA | I-E-I | 1 h, 6 h | Yes | Yes | Constant |
| Shick et al. (1986): fig. 7 | *M. edulis*, Cornwall, England | I-E-I | 8 h | No | Yes | Constant |
| Zenone et al. (2009): abstract | *M. galloprovincialis* and *M. trossulus*, Unspecified | I-E-I | Unspecified | Yes | Unspecified | Variable |
| Connor and Gracey (2012): fig. 1 | *M. californianus*, Malibu, USA | 3 semi-diurnal (I-E-I-E) cycles | 6 h | No | Yes | Constant |
| Peterson et al. (2025): fig. 3 | *M. edulis*, Balgzand, Netherlands | I-E-I-E-I | 0.67 h, ~16 h | No | No | Unspecified |

Our review of previous studies on *Mytilus* mussels also reveals a common experimental setup (Table 1). Most studies have employed a simplistic transition protocol with a single immersion-emersion-reimmersion (I-E-I) sequence, whereas the use of repeated immersion-emersion cycles is rare (Connor and Gracey, 2012). Moreover, few studies have attempted to vary the temperature during emersion (Coleman, 1973; Zenone et al., 2009), despite temperature fluctuations being a defining stressor of intertidal habitats. These limitations can be partly attributed to the lack of laboratory systems capable of simulating a realistic intertidal environment. Using a recently developed intertidal chamber (Pereira et al., 2025), the present study therefore aims to reassess the effects of immersion-emersion cycles and temperature on the heart rate of the blue mussel *M. edulis* under normal conditions. We also report on the responses of mussels following a prolonged emersion event caused by an unexpected chamber malfunction. As part of this study, we introduce a companion software package named rambur to facilitate the design and analysis of intertidal chamber experiments in R (R Core Team, 2026).

## 2 Materials and methods

### 2.1 Animal collection

Blue mussels (*Mytilus edulis*) were collected by hand from a groyne in the Elbe Estuary near Büsum, Germany (54.1247, 8.8713) in May 2025. The mussels were transported in an insulated container to the laboratory on the same day. Upon arrival, the mussels were cleaned of epibionts, and individuals with a shell length of 4.0–5.0 cm were selected for the intertidal chamber experiment.

### 2.2 Intertidal chamber

A detailed introduction to the intertidal chamber (ElectricBlue, Portugal) was given by Pereira et al. (2025). Here, we describe the chamber from a user perspective to facilitate understanding of our experimental setup and subsequent results. Briefly, the chamber contains a lidded experimental tank placed on top of a water storage tank. The experimental tank has drainage holes in its floor to maintain emersion as the default state, and overflow standpipes (11-cm height) to maintain immersion at a constant water level. To establish immersion, a tide pump in the storage tank is activated, continuously transferring water to the experimental tank at a higher flow rate than that of the drainage holes. The water level rapidly rises to the top of the overflow standpipes, which, together with the drainage holes, return excess water to the storage tank. In this two-tank system, the total water volume remains at 42 L, and a partial water change (∼70%) is facilitated through an outlet valve and an inlet pump connected to the storage tank.

Besides immersion status and water change, other environmental variables that can be controlled are light and temperature. Visible light is emitted by dimmable LED strips on the lid of the experimental tank. The lid also has non-luminous heat lamps and circulation fans that increase and homogenize the air temperature during emersion, along with exhaust fans that lower the temperature to the ambient (or room) temperature. Meanwhile, the storage tank is equipped with a submersible heating rod and is connected to an external water chiller via a chiller pump to adjust the water temperature.

The system is governed by an electronic control unit, which receives input from water level and temperature sensors in both tanks and triggers actuators (e.g., pumps and heat lamps) to maintain the target values of environmental variables. The time series of these target values, known as the “experimental profile”, is uploaded to the control unit as a structured text file. At each timestamp, light can be set from 0 to 100% of the maximum brightness (4520 lumens) in 1% increments, and temperature from 0.0 to 60.0 °C in 0.1 °C increments. In contrast, immersion status and water change are binary, with 0 for emersion or no water change, and 1 for immersion or an automatic water change. Between the given timestamps, the control unit applies linear interpolation for temperature and forward-fill step interpolation for all other variables.

### 2.3 Design of experimental profile

First, a diurnal profile with 1-h resolution was constructed to approximate the conditions at the collection site on a typical summer day (Figure 1). Light was modeled as a Gaussian function of time, featuring a 16-h photoperiod (5:00 to 21:00 with at least 1% brightness) and peaking at 13:00 with 100% brightness (Engelmann and Antkowiak, 2016). Terrestrial air temperature and offshore surface water temperature were modeled as sinusoidal functions, both peaking at 15:00 to mimic a thermal lag following the light peak (Chow and Levermore, 2007). Based on summer 2024 data from the nearby Elpersbüttel weather station (54.0691, 9.0105; Deutscher Wetterdienst, 2024) and the ElbeWR buoy (53.9965, 8.1137; Bundesamt für Seeschifffahrt und Hydrographie, 2025), the air temperature was set to a mean of 17 °C with a range of 8 °C, while the water temperature was set to a mean of 19 °C with a range of 1 °C. A symmetric, semi-diurnal immersion-emersion cycle was simulated under a simplified 24-h tidal day (Connor and Gracey, 2012). Emersion was set from 0:00 to 6:00 and 12:00 to 18:00, while immersion was set from 6:00 to 12:00 and 18:00 to 0:00. The target exposure temperature for the experimental tank was then determined by assuming that the intertidal zone temperature matches the terrestrial air temperature during emersion and the offshore surface water temperature during immersion. Due to manufacturer-noted technical risks, the automatic water change was disabled (i.e., maintained at 0).

**Figure 1.**
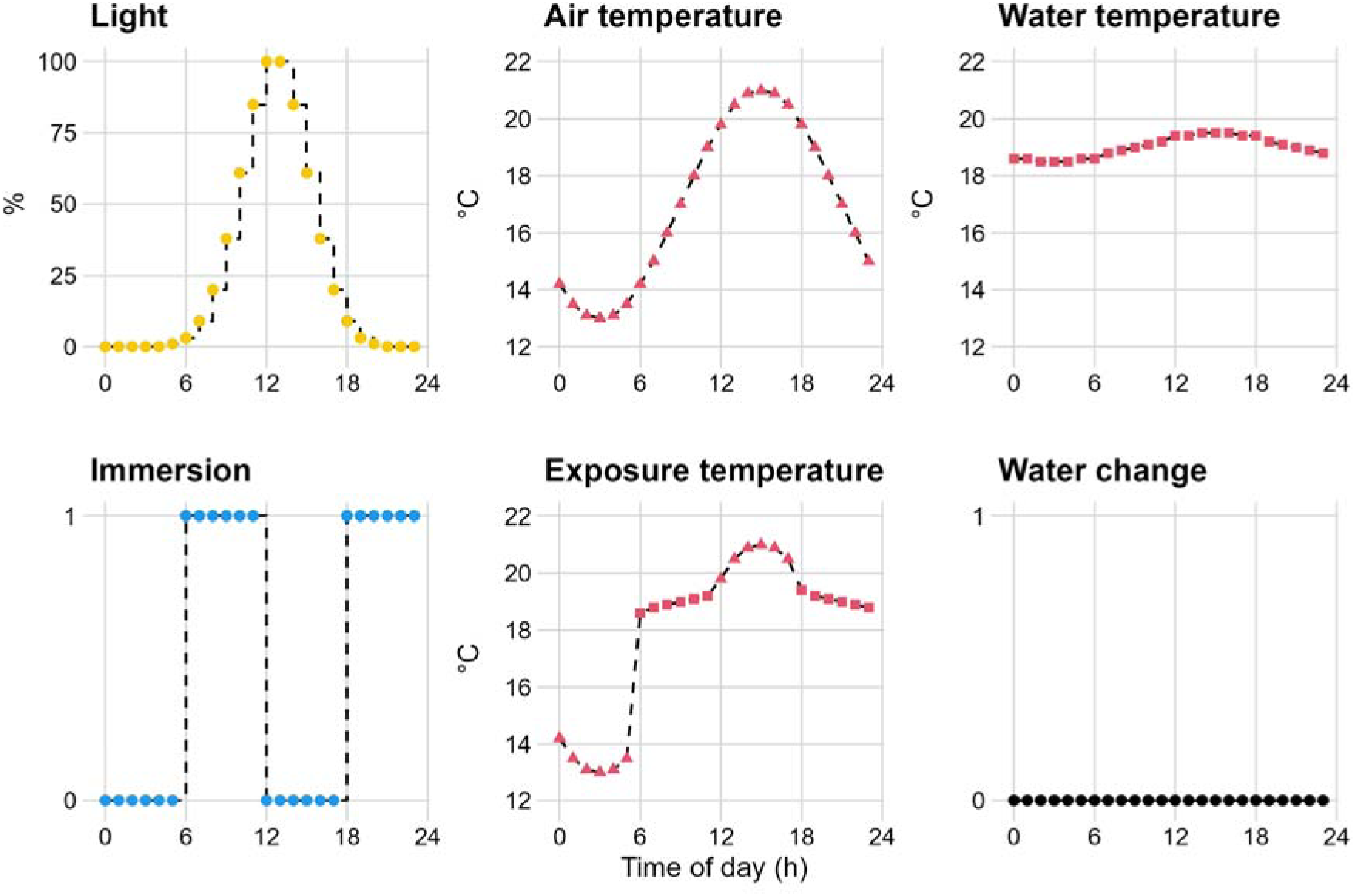
Diurnal profile of light, temperature, immersion status, and water change with 1-h resolution. The target exposure temperature was derived by first simulating the air and water temperatures separately, then selecting between them according to the immersion status. Dashed lines indicate the expected interpolation between hourly timestamps by the intertidal chamber.

Finally, to simulate a consistent environment for assessing long-term physiological responses, the diurnal profile was repeated identically for one month. This multiday experimental profile was then exported to a text file ready for chamber operation. The entire process was performed using the chamber.design() function from the rambur package (Table 2).

**Table 2:** Main functions from the rambur package for experimental design and analysis.

| Function | Description |
| --- | --- |
| `chamber.design()` | Design of experimental profile for intertidal chamber |
| `chamber.read()` | Read log files from a single chamber |
| `chamber.check()` | Check chamber conditions against profile |
| `robos.read()` | Read log files from multiple robomussels |
| `pulse.read()` | Read log files from a single Pulse logger |
| `pulse.extract()` | Extract heart rate from Pulse logger records |

### 2.4 Chamber operation

The experimental profile was uploaded to two identical intertidal chambers, designated as Chamber A and Chamber B, each housing 200 mussels. The chambers were operated at room temperature and filled with artificial seawater at salinity 30 (Pro-Reef, Tropic Marin, Germany). Water changes were performed twice a week during the afternoon emersion by manually triggering the outlet valve and the inlet pump. After each water change, 10 mL of a live microalgal blend (Live Marine Phytoplankton–Premium Reef Blend, Sustainable Aquatics, USA) was added to each chamber.

Throughout the operation, data from the chamber sensors and actuators were recorded by the control units every 10 s and stored in hourly CSV log files. The files from each chamber were imported into R using the chamber.read() function. A visual comparison between the designed experimental profile and the actual chamber conditions was conducted using the chamber.check() function.

Alongside the mussels, two biomimetic temperature loggers, or “robomussels” (EnvLogger T2.4, ElectricBlue, Portugal; Pereira et al., 2026), were deployed in each chamber to estimate body temperature. The temperature was recorded every 15 min and stored in a CSV log file. The files from the two robomussels were imported using the robos.read() function.

### 2.5 Heart rate monitoring

Ten mussels per chamber, labeled A1 to A10 in Chamber A and B1 to B10 in Chamber B, were randomly selected for continuous heart rate monitoring for 26 days with two 10-channel Pulse V2 loggers (ElectricBlue, Portugal; Lima et al., 2025). Briefly, a wired photoplethysmography (PPG) sensor was attached to the shell of each mussel above its heart using a liquid contact adhesive (UHU, Germany). The sensor emitted infrared light and detected the backscattered light to track volumetric changes in the circulatory structures associated with each heartbeat. This optical signal was converted into an analog electrical signal, which was then digitized at a sampling rate of 5 Hz (one sample every 0.2 s) and stored in hourly CSV log files. The files from each Pulse logger were imported into R using the pulse.read() function.

During the experiment, if a mussel died, the sensor was redeployed to a randomly chosen replacement mussel from the same chamber.

### 2.6 Heart rate extraction

An autocorrelation-based algorithm was developed to extract heart rate from the PPG signal. Specifically, the Pulse logger records were divided into non-overlapping 1-min (300-sample) windows (Figure 2). For each window, the autocorrelation function (ACF) was computed to assess the similarity between the signal and its lagged versions (Venables and Ripley, 2002). As markers of relatively high similarity, all ACF peaks (local maxima) at non-zero integer lags were identified. Quadratic interpolation was then applied to estimate the true peaks with more precise fractional lags and correlation values (Smith, 2011).

**Figure 2.**
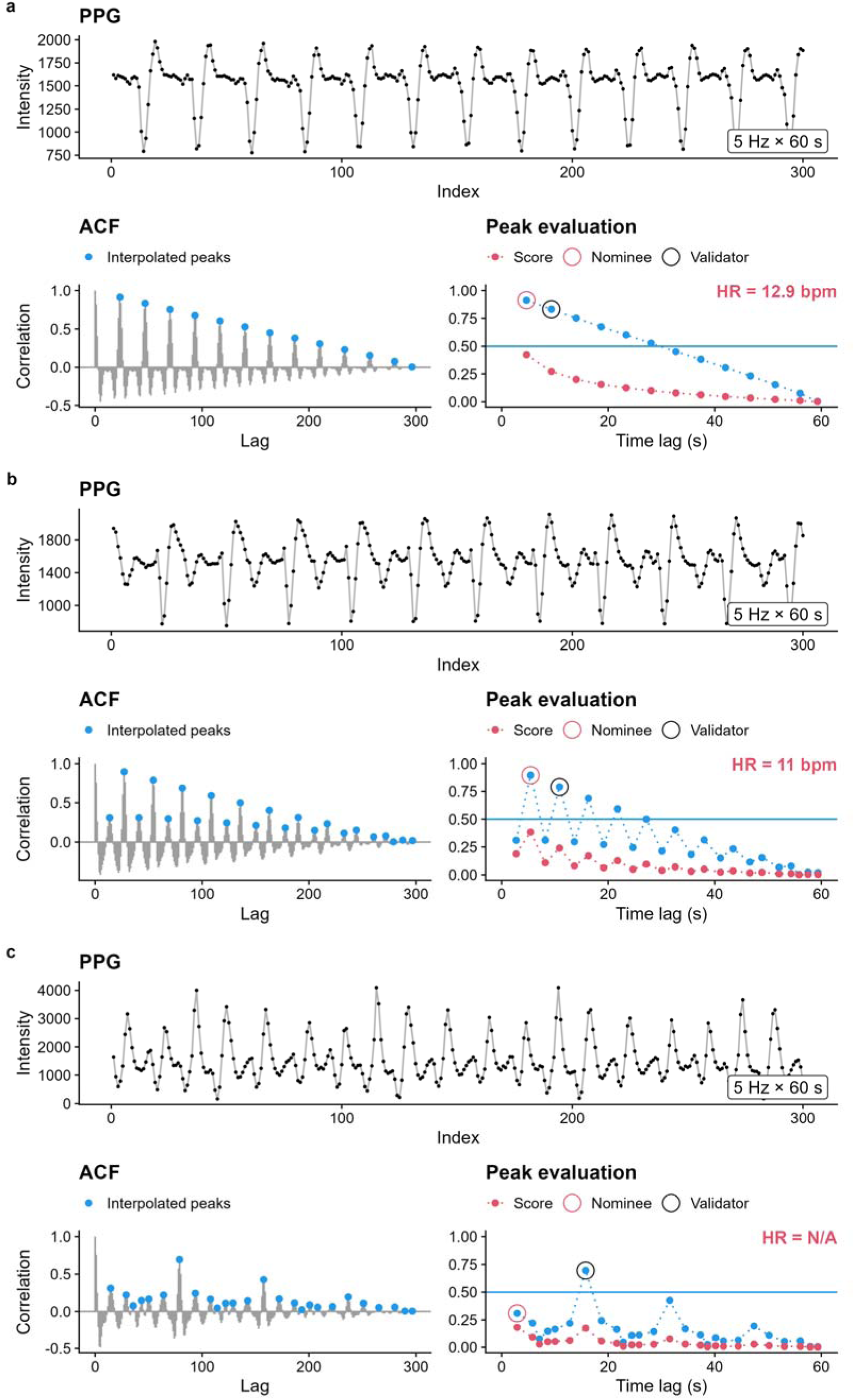
Three examples of heart rate extraction from photoplethysmography (PPG) signals. Each example shows a 1-min PPG window with 300 digital samples connected by lines to visually reconstruct the analog signal (top), the autocorrelation function (ACF) across integer lags and the interpolated peaks at fractional lags (bottom left), and the search for the prominent peak along with the extracted heart rate (HR) (bottom right). The first example (a) shows a regular signal where the prominent peak is both the earliest and the highest among the interpolated peaks. The second example (b) shows a signal with dicrotic pulses. Here, the prominent peak is the highest, but not the earliest. The third example (c) shows an irregular signal with variable beat morphology and a slower natural rhythm (a stronger pulse every 5 beats). In this case, the prominent peak is the earliest, but not the highest. Power-law scoring with the parameter = *0.5* successfully nominates the prominent peak in all examples (peaks and scores connected by dashed lines for visual clarity). In the first two cases, the nominees pass all quality checks, including evaluation against a subsequent validator. However, the nominee in the last case fails the first check by being lower than the minimum threshold = *0.5* (horizontal blue line), yielding an indeterminate heart rate (N/A).

Typically, shifting the signal by its fundamental period, i.e., one interbeat interval, produced a prominent peak in the ACF, while shifting by multiples of the interbeat interval resulted in subsequent decaying peaks (Figure 2a). Accurately identifying this prominent peak is key to determining the interbeat interval and the corresponding heart rate.

A naive choice for the prominent peak would be the earliest or the highest among the interpolated peaks. However, the earliest peak in the ACF may result from complex intrabeat features and correspond to only a fraction of the true interbeat interval (Figure 2b). Using this early peak would lead to an overestimation of heart rate (a harmonic error). Likewise, signal artifacts and natural rhythms can cause the highest peak to occur at a lag corresponding to a multiple of the true interbeat interval (Figure 2c). Using this late peak would result in an underestimation of heart rate (a sub-harmonic error).

To avoid these errors, a two-step procedure involving nomination and validation was implemented. In the nomination step, all peaks were scored using a power-law function:

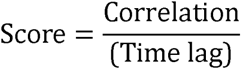

with a non-negative parameter . The higher the value of, the greater the penalty for later peaks. The two highest-scoring peaks were then identified, with the earlier peak acting as the nominee for the prominent peak, and the later peak serving as the validator.

In the validation step, the nominee was subjected to multiple quality checks. First, its correlation value had to exceed a minimum threshold to indicate strong periodicity in the PPG signal. Next, the correlation value of the nominee had to be higher than that of the validator, and the validator had to occur at twice the nominee’s time lag, confirming the presence of an echoing, decaying peak. Lastly, the nominee had to be preceded by at least one negative correlation value as evidence of a genuine oscillatory pattern. If the nominee passed all the checks, its corresponding interbeat interval was used to calculate the heart rate (in beats per minute, bpm) (Figure 2a, b). Otherwise, the algorithm returned an indeterminate heart rate (N/A) for that window (Figure 2c). This often occurred during periods of irregular heartbeats (i.e., inconsistent interbeat intervals) or very slow to absent heart rates (fewer than two beats in the window, meaning an echoing peak could not exist).

The described algorithm was executed using the pulse.extract() function. The optimal values for and were determined using a grid search approach, with ranging from 0.0 to 2.0 in 0.5 increments and ranging from 0.0 to 0.9 in 0.1 increments. Heart rates were manually counted for a 2-h subset of records and compared with the rates generated by the algorithm across all combinations of and . The values that minimized error and maximized coverage were found to be = *0.5* and = *0.5*. These values were then applied to all records to extract heart rates.

### 2.7 Statistical analyses

First, the 10-s resolution environmental data were averaged to a 1-min resolution to match the heart rate data. This preprocessing step was handled by the chamber.read() function. As detailed in the Results section, an unexpected malfunction occurred in Chamber B, causing a prolonged emersion event and increased mortality. Given the diverging environmental conditions, we separated the analyses of heart rate in the two chambers.

#### 2.7.1 Chamber A

Heart rate was analyzed using a generalized additive mixed model (GAMM). The model included immersion status (emersion versus immersion) as a binary predictor. Conditioned on this predictor, smooth functions were applied to the time elapsed since the last transition to model non-linear heart rate dynamics. This elapsed time was log- transformed to better resolve changes in heart rate immediately following transitions. Similarly, conditioned on immersion status, smooth functions were applied to temperature to model the thermal responses of heart rate. Because temperature and light were highly correlated by design, light was not included in the model. To account for individual variations in the longitudinal trajectories of heart rate, mussel-specific smooth functions were applied to the time elapsed since the start of the experiment. Additionally, mussel volume was estimated from measured shell length, height, and width assuming an ellipsoid shape, and this size metric was included as a continuous covariate. Food supply was also added as a binary covariate, effective during the immediate immersion and subsequent emersion periods following the manual water changes and microalgal addition.

Following the specification of predictors, the distribution of the response variable (heart rate) was evaluated. Three candidate models using Gaussian, scaled t (scat), and gamma distributions, all with a log link function, were fitted to a random 1% subset of the data using maximum likelihood (ML) estimation. Based on the Akaike Information Criterion (AIC), the scaled t-distribution provided the best fit, likely because of the heavy-tailed heart rate data. Using the scaled t-distribution and fast restricted maximum likelihood (fREML), a naive model was fitted to the full dataset, and the residuals were extracted to estimate the first-order autoregressive (AR1) parameter (=0.87). This parameter reflected the temporal autocorrelation of heart rate within individual mussels and was subsequently used to refit the final model.

Given that heart rate values were missing when the PPG signal failed the quality checks of the extraction algorithm, we additionally examined whether the missingness could be explained by the observed variables. Specifically, heart rate extraction success was coded as a binary response and analyzed using a GAMM with a binomial distribution, a logit link function, and an AR1 correlation structure (estimated = *0.67*). The predictors of this auxiliary model remained the same as in the main model.

#### 2.7.2 Chamber B

Data recorded during the prolonged emersion event were excluded from the analysis due to the extreme environmental conditions that were not comparable to those of the rest of the experiment. The focus here was on comparing heart rate patterns before and after the event. Specifically, heart rate was analyzed using a hierarchical generalized additive model (HGAM; Pedersen et al., 2019) with a scaled t-distribution, a log link function, and an AR1 correlation structure (estimated = *0.84*). The model included experimental phase (pre- versus post-event) and immersion status as two binary predictors, along with their interaction. Conditioned on this interaction, smooth functions were applied to the log-transformed elapsed time since the last transition and to temperature. As some originally monitored mussels died during the event and replacement mussels were monitored post-event, the longitudinal heart rate trajectories were modeled using a hierarchical structure with three components: global smooth functions applied to the elapsed time since the start of each experimental phase, random intercepts for individual mussels, and mussel- and phase-specific smooth functions applied to the same elapsed time variable. Given the negligible effects of mussel volume and food supply on heart rate in Chamber A (see the Results section), these covariates were omitted.

Finally, the partial effects of the predictors in the GAMMs and HGAM were assessed via conditional predictions (Heiss, 2022). Briefly, the expected values of the responses were estimated by evaluating the focal predictors across their observed ranges while holding the remaining variables constant at their representative levels. Here, mussel-specific smooths and random intercepts were excluded to estimate population-level effects and were retained only for individual-level longitudinal trajectories.

All described analyses were conducted in R v4.6.1 (R Core Team, 2026). The GAMMs and HGAM were fitted using the mgcv package (Wood et al., 2017), and conditional predictions were obtained using the marginaleffects package (Arel-Bundock et al., 2024).

## 3 Results

### 3.1 Chamber conditions and mortality

Throughout the 26-day experiment, the actual conditions in Chamber A generally matched the designed experimental profile (Figure 3a). There was virtually no discrepancy in light output, aside from brief interruptions during manual water changes when the automatic system was temporarily disabled (Figure 3b). The tide pump also reliably followed the programmed immersion-emersion cycles. However, while the water filled the experimental tank rapidly when the pump turned on, it drained gradually when the pump turned off, causing the actual immersion periods to last slightly longer than the intended 6 h. Regarding thermal conditions, the actual temperature closely matched the design during immersion but was substantially higher during emersion, particularly at night. This deviation was due to the limited air cooling capacity of the chamber, which was constrained by the room temperature. Additionally, the body temperature recorded by the robomussels closely tracked the water temperature during immersion, but remained consistently higher than the air temperature during emersion.

**Figure 3.**
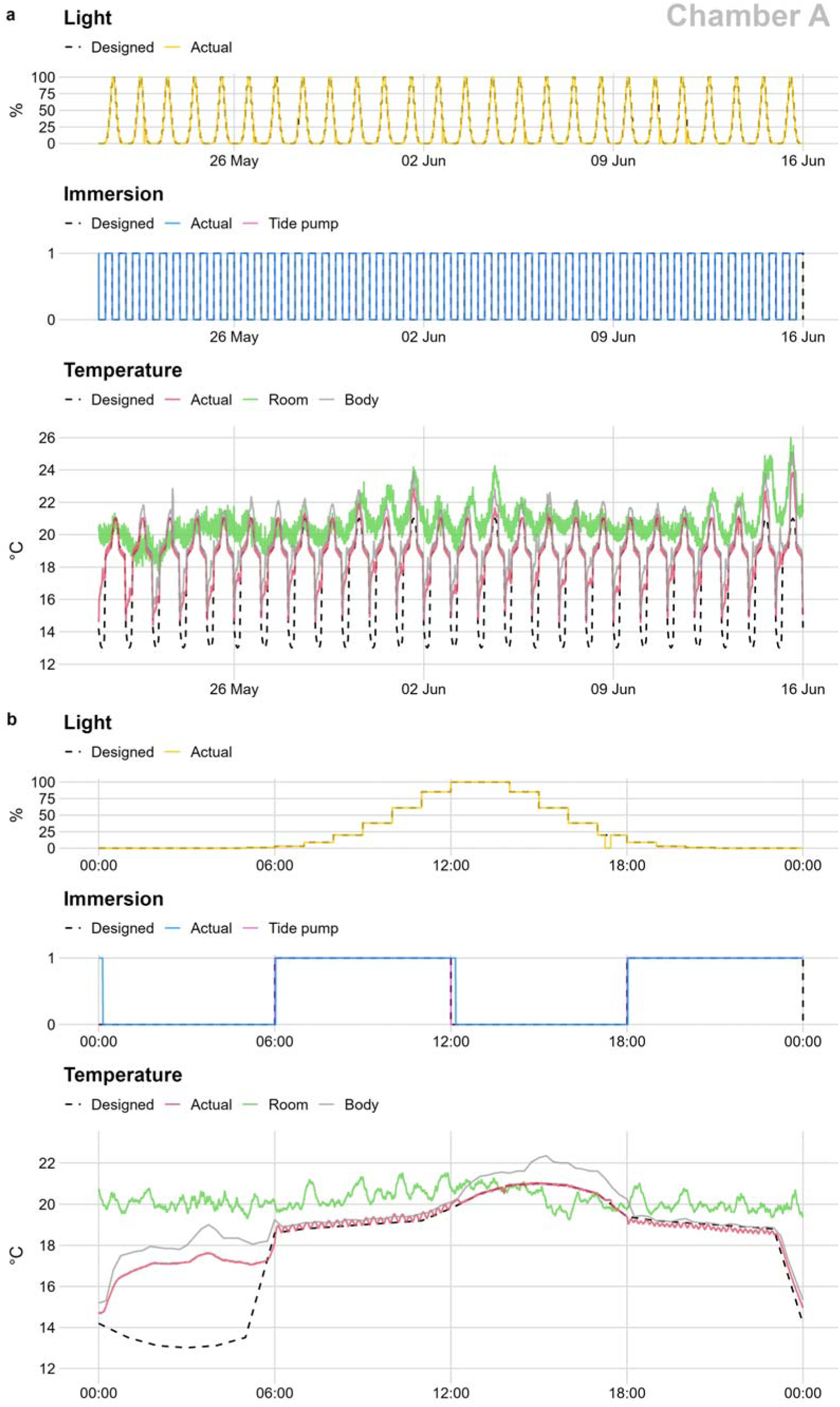
Overview of environmental conditions in Chamber A throughout the experiment (21 May to 15 June) (a) and a closer look at conditions on a typical day (26 May) (b). The actual conditions of light (LED brightness), immersion status (average of three sensors), and temperature (average of three sensors) are compared with the designed experimental profile. The tide pump status, room temperature, and body temperature (average of two robomussels) are also provided.

The observations described for Chamber A largely held true for Chamber B, with the exception of two malfunctions (Figure 4a). These occurred on the afternoons of 28 May and 6 June, when the outlet valve erroneously opened, causing substantial water loss from the chamber. The first malfunction triggered an early immersion, despite the tide pump falsely recording as inactive (Figure 4b). This immersion was only partial, as indicated by the increased temperature driven by the ambient air. The system was then promptly rebooted to resume the experiment. The second malfunction occurred similarly but went undetected over a holiday period until 10 June, resulting in a prolonged emersion event with elevated temperatures (Figure 4c).

**Figure 4.**
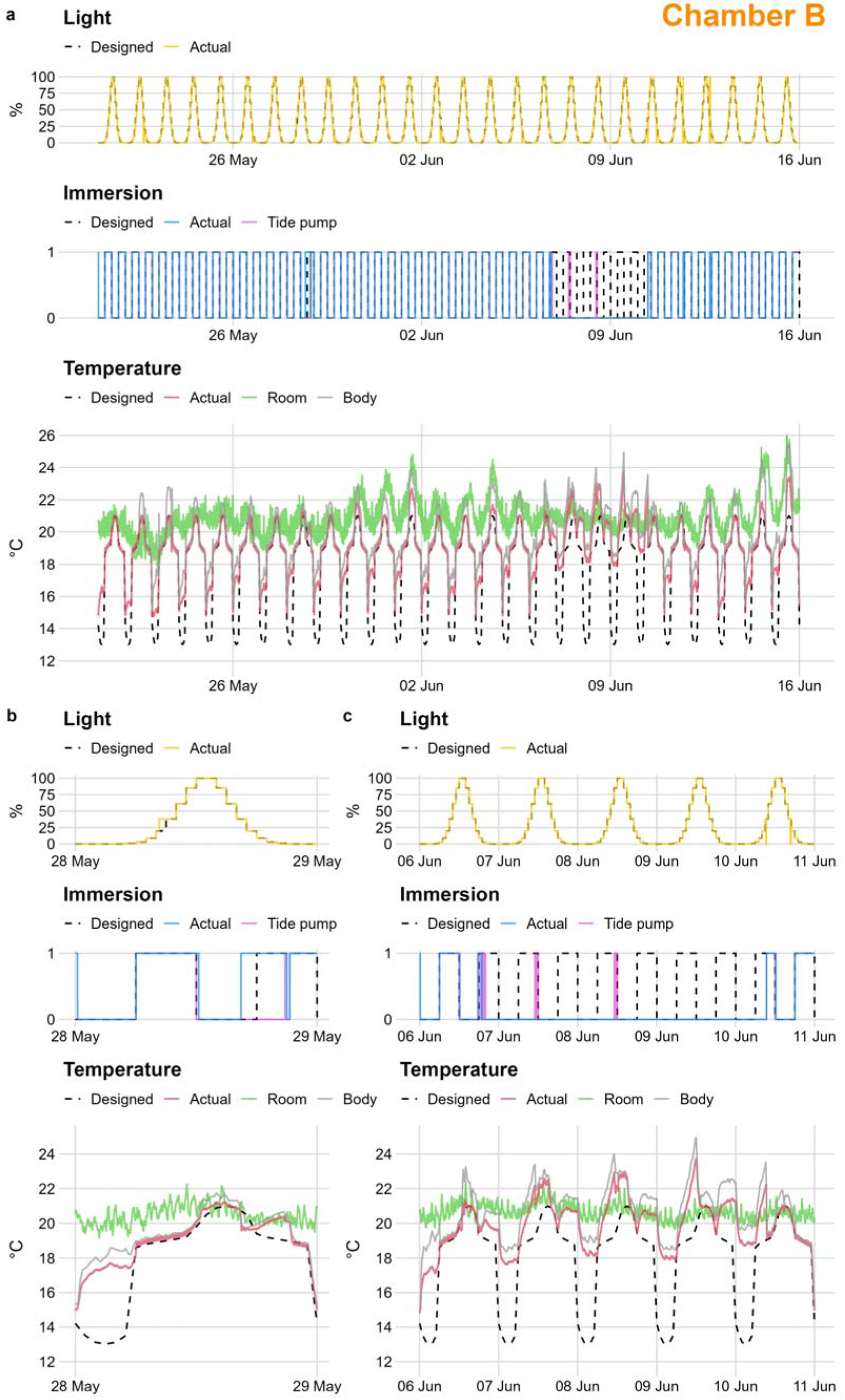
Overview of environmental conditions in Chamber B throughout the experiment (21 May to 15 June) (a) and a closer look at two malfunctions on 28 May (b) and 6 June (c). The actual conditions of light (LED brightness), immersion status (average of three sensors), and temperature (average of three sensors) are compared with the designed experimental profile. The tide pump status, room temperature, and body temperature (average of two robomussels) are also provided.

This extreme event was associated with increased mortality in Chamber B upon detection and in the following days, whereas mortality in Chamber A remained consistently low (Figure 5). Among the deaths in Chamber B on 10 and 11 June were mussels B2 and B7, which were replaced by new mussels (B2R and B7R) for post-event heart rate monitoring. In Chamber A, mussel A8 died on 6 June without an apparent environmental cause, suggesting an underlying health issue. Therefore, its heart rate data were excluded from the analyses. Data from its replacement mussel were also discarded due to the short temporal coverage.

**Figure 5.**
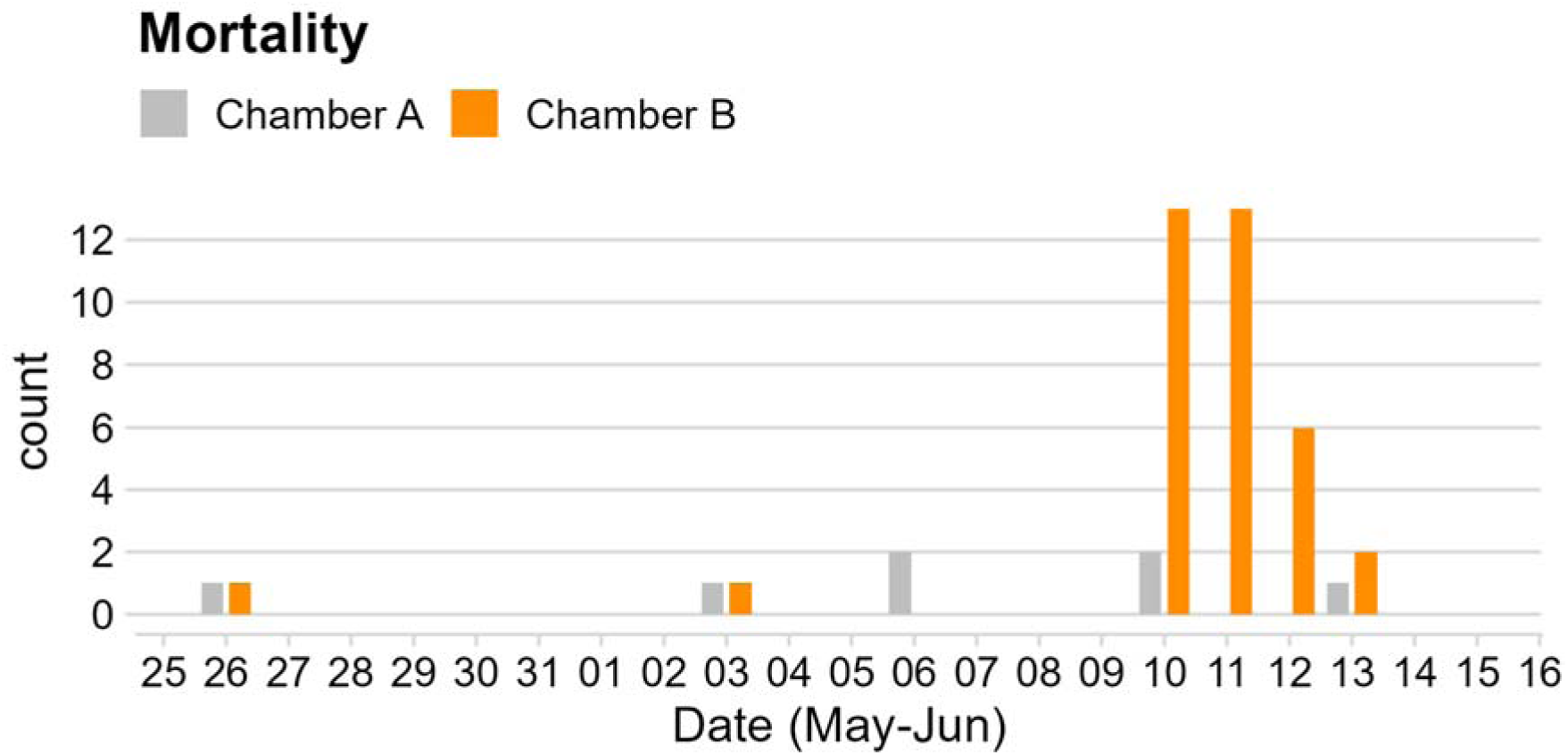
Mortality recorded in each chamber out of the initial 200 mussels, from the first death (26 May) to the end of the experiment (15 June). Note that the zero counts between 7 and 9 June indicate a lack of observations during the holiday period.

### 3.2 Heart rate extraction success

The auxiliary GAMM for Chamber A explained 35.0% of the total deviance in heart rate extraction success (n = 336 609). The success probability showed opposing trends depending on the direction of the transition (Figure 6a). Upon transition to emersion, the probability declined from above 60% to 6% in less than 2 h and stayed below 20% for the remainder of emersion. In contrast, transitioning to immersion increased the probability from below 25% to above 70% within 20 min, a level maintained throughout the rest of immersion.

**Figure 6.**
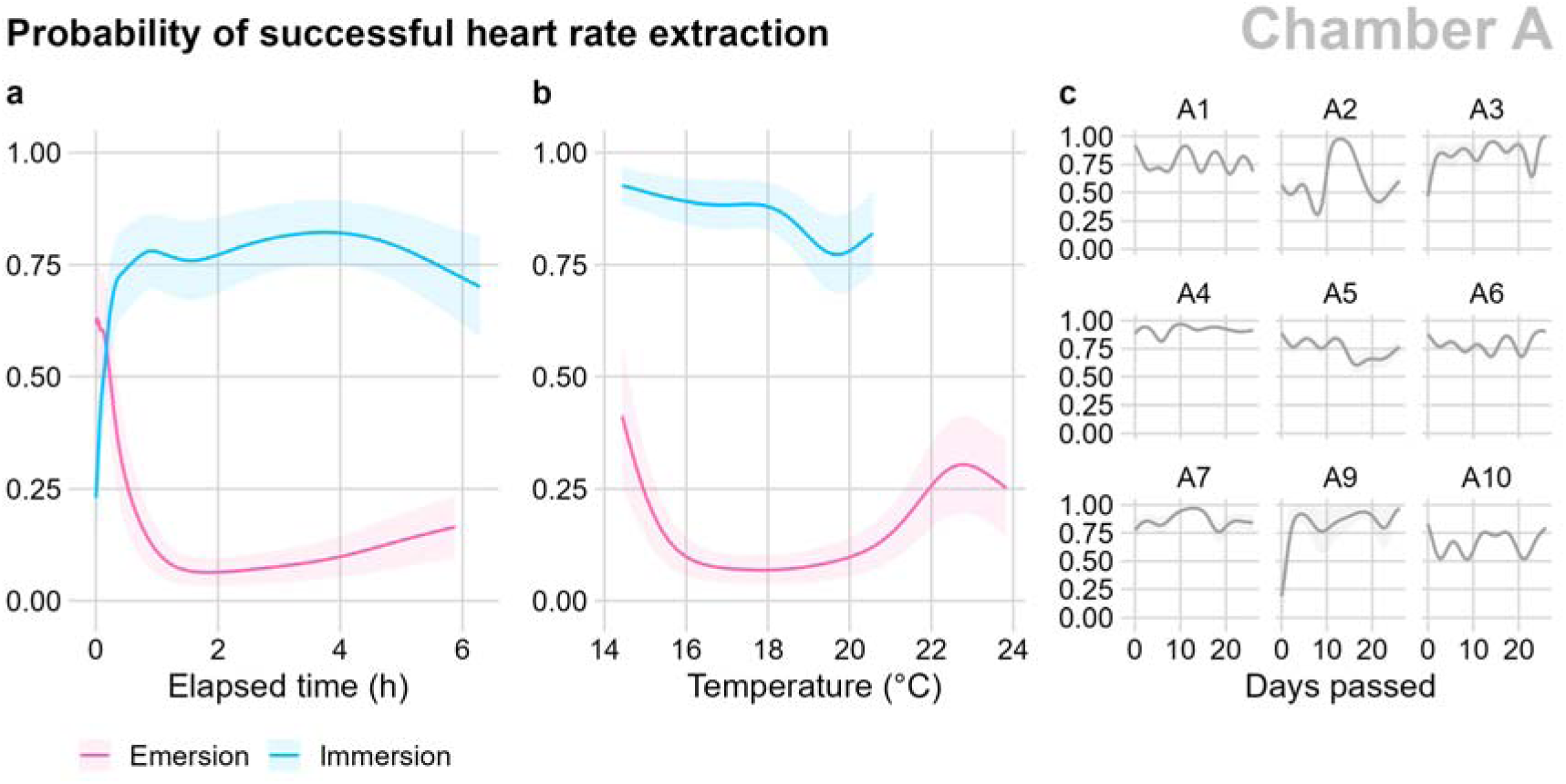
Conditional predictions of the probability of successful heart rate extraction in Chamber A. Focal predictors include immersion status and elapsed time since the last transition (a), immersion status and temperature (b), and individual mussels and elapsed time since the start of the experiment (c). Mussel-specific smooths are excluded in (a) and (b) to estimate population-level effects. When not acting as focal predictors, variables are held constant at representative levels: immersion status set to immersion, elapsed time of 3 h since the last transition, temperature of 19 °C, elapsed time of 13 days since the start of the experiment, mussel volume of 10 mL, and no food supply. Shaded areas denote 95% confidence intervals.

The effects of temperature on extraction success also varied by immersion status (Figure 6b). During immersion, the success probability remained relatively stable above 75% across the temperature range. During emersion, the probability was below 10% at intermediate temperatures but increased at both higher and lower temperatures.

Longitudinally, the probability of successful heart rate extraction in individual mussels fluctuated during the experiment but exhibited no clear directional trend (Figure 6c). Regarding the covariates, for each 1 mL increase in mussel volume, the odds of extraction success decreased by 25.4% (*p*-value = 0.02), whereas the odds increased by 2.3% with food supply (*p*-value = 0.49).

### 3.3 Heart rate dynamics following transitions

The main GAMM for Chamber A and the HGAM for Chamber B explained 56.6% and 62.2% of the total deviance in heart rate, respectively (n = 152 754 and 154 955). Heart rate was highly dependent on immersion status, with distinct non-linear dynamics following transitions (Figures 7a and 8a). In Chamber A (Figure 7a), heart rate increased slightly during the first 10 min of emersion, from nearly 21 bpm to almost 22 bpm. For the next 10 min, it decreased slightly but remained above 20 bpm, before declining sharply and stabilizing at a low baseline of around 6 bpm for the remainder of emersion. The transition to immersion triggered an immediate increase in heart rate, peaking at nearly 26 bpm within the first few minutes, then fluctuating briefly before gradually decreasing within 2 h to a high baseline of around 20 bpm.

**Figure 7.**
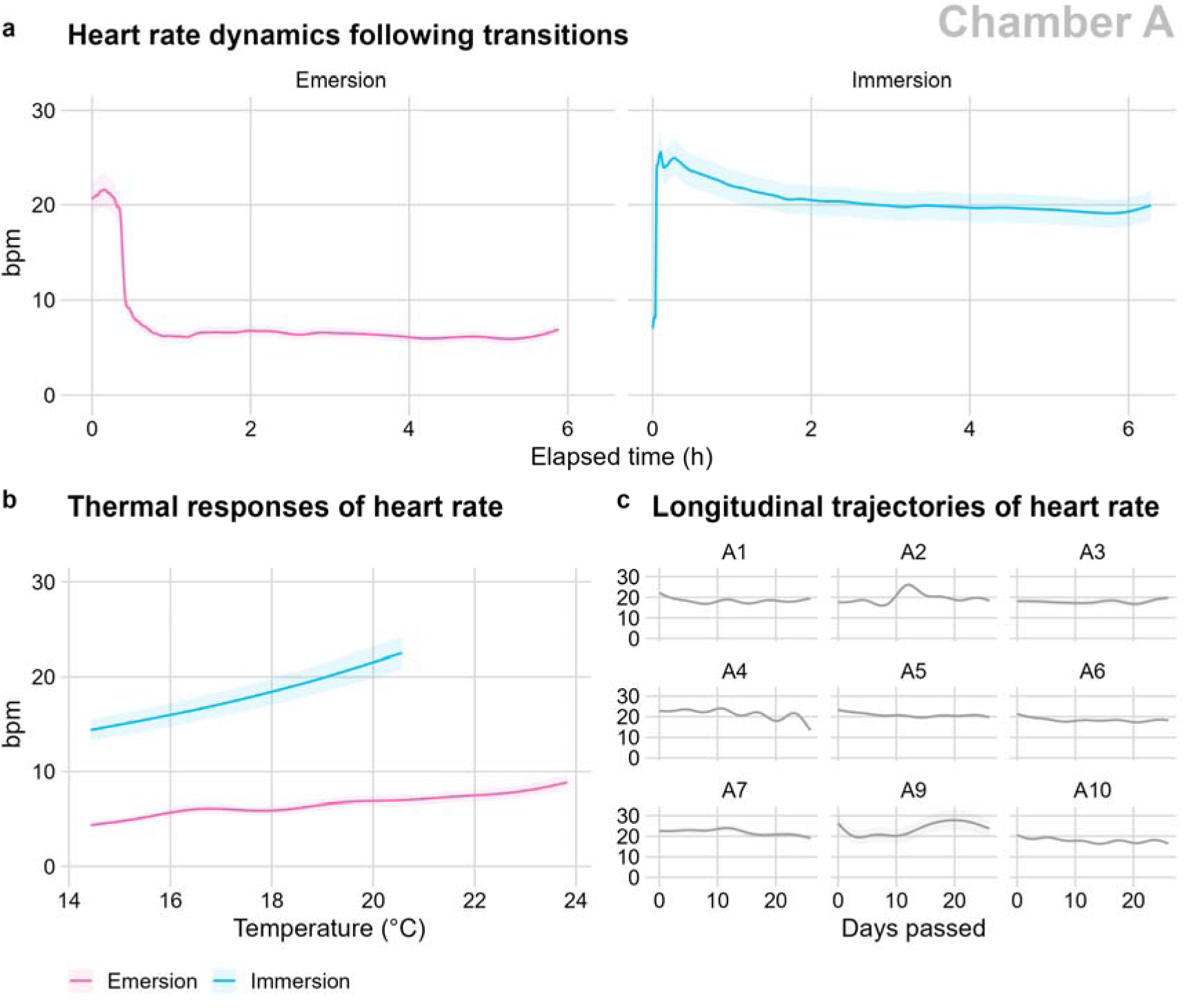
Conditional predictions of heart rate in Chamber A. Focal predictors include immersion status and elapsed time since the last transition (a), immersion status and temperature (b), and individual mussels and elapsed time since the start of the experiment (c). Mussel-specific smooths are excluded in (a) and (b) to estimate population-level effects. When not acting as focal predictors, variables are held constant at representative levels: immersion status set to immersion, elapsed time of 3 h since the last transition, temperature of 19 °C, elapsed time of 13 days since the start of the experiment, mussel volume of 10 mL, and no food supply. Shaded areas denote 95% confidence intervals.

**Figure 8.**
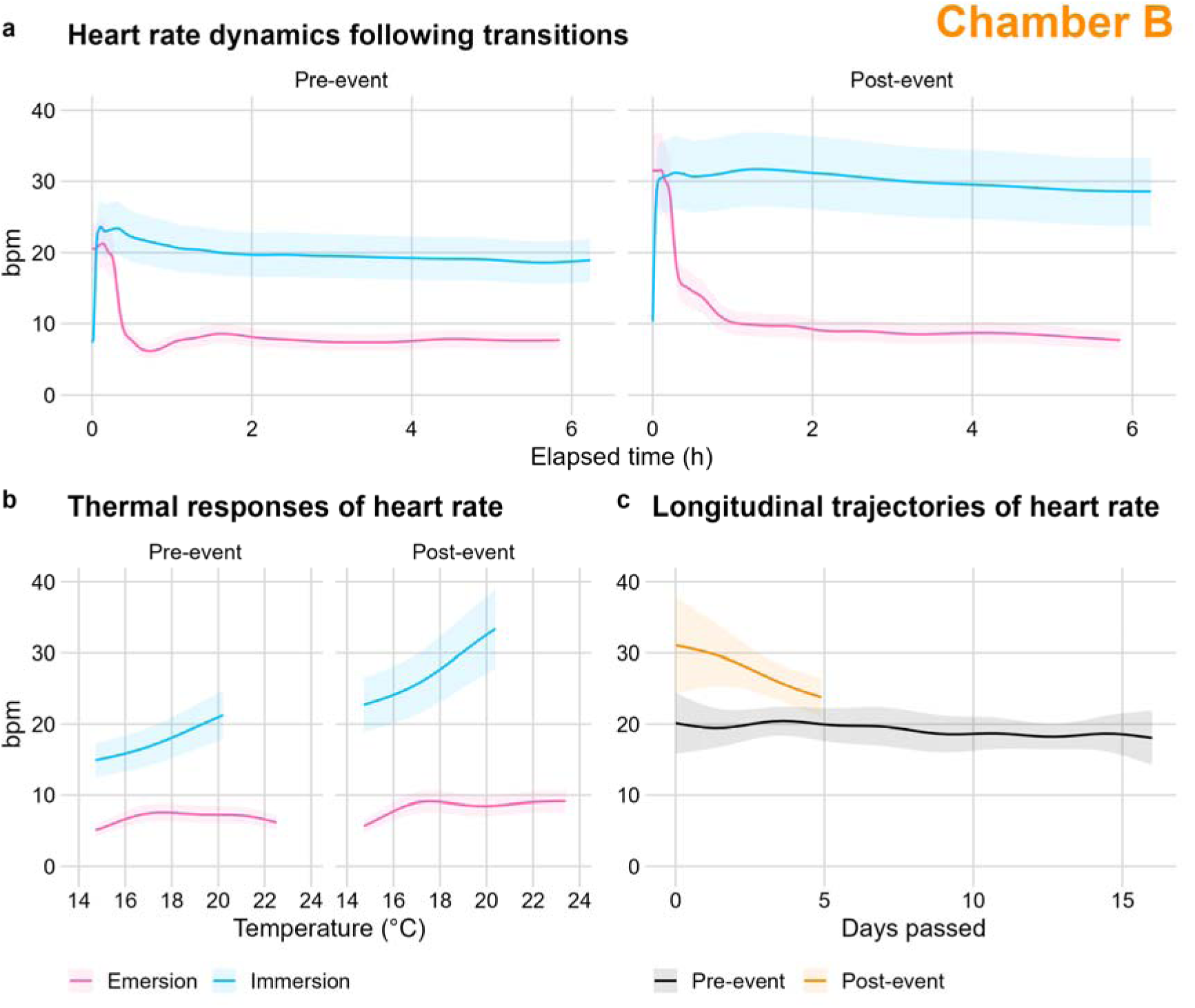
Conditional predictions of heart rate in Chamber B. Focal predictors include experimental phase, immersion status, and elapsed time since the last transition (a); experimental phase, immersion status, and temperature (b); and experimental phase and elapsed time since the start of each phase (c). Mussel-specific smooths and random intercepts are excluded to estimate population-level effects. When not acting as focal predictors, variables are held constant at representative levels: immersion status set to immersion, elapsed time of 3 h since the last transition, temperature of 19 °C, and elapsed time of 1 day since the start of each experimental phase. Shaded areas denote 95% confidence intervals.

In Chamber B (Figure 8a), heart rate dynamics in the pre-event phase were similar to those observed in Chamber A. Specifically, the onset of emersion was characterized by a slight increase in heart rate, from above 20 bpm to over 21 bpm, followed by a decline to a low baseline of around 8 bpm. The transition to immersion also induced an immediate increase in heart rate to nearly 24 bpm, followed by a brief fluctuation and a gradual return to a high baseline of around 19 bpm. In contrast, heart rate dynamics differed markedly in the post-event phase. Here, the onset of emersion featured a brief plateau in heart rate above 31 bpm lasting less than 10 min, followed by a decline to a low baseline of around 9 bpm. Upon immersion, heart rate increased immediately to above 30 bpm and remained elevated at 29–31 bpm for the rest of immersion.

### 3.4 Thermal responses of heart rate

The thermal responses of heart rate were also affected by immersion status and experimental phase (Figures 7b and 8b). During immersion, heart rate in Chamber A (Figure 7b) showed a positive, monotonic relationship with temperature, with an average increase of 1.3 bpm per °C. Similarly, pre-event heart rate in Chamber B (Figure 8b) recorded an average increase of 1.2 bpm per °C. In comparison, the post-event average increase was higher, reaching 1.9 bpm per °C. During emersion, heart rate exhibited weak thermal responses across chambers and experimental phases, with average increases of 0.3–0.5 bpm per °C.

### 3.5 Longitudinal trajectories of heart rate and effects of covariates

During the 26-day experiment in Chamber A (Figure 7c), the longitudinal trajectories of heart rate across individual mussels were relatively stable, remaining largely between 18 and 21 bpm with no consistent long-term increases or decreases. Similarly, the pre-event population-level trajectory in Chamber B remained stable at 19–20 bpm (Figure 8c). In comparison, post-event heart rate started higher at 31 bpm, then declined progressively over the subsequent five days, dropping below 24 bpm by the end of the experiment.

Furthermore, within the main GAMM for Chamber A, the effects of mussel volume and food supply on heart rate were negligible. As mussel volume increased by 1 mL, the heart rate decreased by 1.9% (*p*-value = 0.32), and when food was supplied, the heart rate decreased by 0.3% (*p*-value = 0.25).

## 4 Discussion

### 4.1 Intertidal simulation

In this study, we used a mechanistic approach to design the experimental profile (Figure 1). Here, environmental variables were constructed with idealized patterns, e.g., sinusoidal temperature curves and a symmetric immersion-emersion cycle. We also assumed that the intertidal zone temperature could be expressed as a simple function of the terrestrial air temperature, offshore surface water temperature, and immersion status. This approach can be refined to more accurately simulate field conditions. For example, an asymmetric cycle with an extended emersion period can be used for mussels originating from the high intertidal zone (Leeuwis and Gamperl, 2022). Additionally, the function used to derive the intertidal zone temperature could be adjusted in the future when *in situ* temperature data from the site become available (Pereira et al., 2025).

The implementation of the designed profile also revealed several issues with the intertidal chambers. First, the control unit applied linear interpolation exclusively to temperature, which could cause noticeable thermal ramps at the end of immersion and emersion (Figure 3b). Specifically, between 5:00 and 6:00, the air temperature was forced to increase to match the high target water temperature at the onset of the next immersion. Similarly, between 23:00 and 0:00, the water temperature was forced to decrease to match the low target air temperature at the beginning of the next emersion. These thermal ramps could hypothetically occur in the field when a flood tide of warmer water heats the air above the intertidal zone at the end of emersion, or when an ebb tide leaves a thin layer of water at the end of immersion that is subsequently cooled by the colder air above it. However, the extent to which this holds true in the field requires further validation using *in situ* data. In the meantime, a technical workaround to prevent these ramps is to design the profile with a much finer resolution, e.g., 5 min instead of 1 h. Because the hardware might not heat or cool rapidly enough to complete a steep ramp within 5 min, the temperature would remain relatively close to its preceding target value until the transition actually occurs.

Second, the chambers lacked active air cooling and relied solely on the exhaust fans to lower the air temperature during emersion, which proved ineffective given the room temperature (Figure 3). To accurately implement the designed profile, the chambers would need to be operated inside a cold room. However, this solution introduces operational risks for the water chiller, which is not designed to work at low ambient temperatures. One alternative would be to decommission the chiller and allow the cold room to passively cool the water, albeit at the expense of speed and precision.

Finally, the two malfunctions of the outlet valve in Chamber B suggest that this component is currently unreliable, despite its critical role in the continuous operation of the chamber (Figure 4). As a quick fix, an additional manual valve can be installed to block the outlet and prevent water loss in the event of a malfunction.

### 4.2 Heart rate under normal conditions

The current study confirmed the presence of an emersion spike in the heart rate of the blue mussel *Mytilus edulis*. In both Chamber A (Figure 7a) and Chamber B before the extreme event (Figure 8a), the spike occurred within 10 min of emersion, with a modest increase of about 1 bpm. This finding is consistent with observations in the California mussel *M. californianus*, where the spike occurred within 5 min of emersion with an increase of almost 4 bpm (Bayne et al., 1976). Emersion spikes were also reported in the Mediterranean mussel *M. galloprovincialis* and the bay mussel *M. trossulus* (Zenone et al., 2009), although their timing and magnitude were not specified. Furthermore, the emersion spike is not unique to *Mytilus* mussels. In *Cerastoderma* cockles, the spike occurred within 5 min of emersion with an increase of 2 bpm (Boyden, 1972). Similar spikes were also observed in *Crassostrea* oysters, with an increase of less than 4 bpm within a few minutes (de Carvalho et al., 2022, 2026). Taken together, these results suggest that emersion spikes in intertidal bivalves are inherently small and short-lived. This subtle nature probably explains why they have been overlooked in previous studies (Table 1).

Mechanistically, the emersion spike likely represents an acute compensatory response to internal hypoxia. Upon emersion, the animals close their valves to avoid desiccation, but this closure simultaneously halts the oxygen supply (Helm and Trueman, 1967). The brief increase in heart rate may therefore facilitate the utilization of residual oxygen retained within their mantle cavity (de Carvalho et al., 2022). This mechanism is supported by classic studies on the effects of aquatic hypoxia on the heart rate of *M. edulis*. Specifically, mussels in stagnant, sealed water (Schlieper, 1955) and those in water subjected to progressive hypoxia via nitrogen bubbling (Bayne, 1971) both showed brief increases in heart rate prior to an eventual decline.

The low heart rate baseline observed following the emersion spike (Figures 7a and 8a) is expected, given the well-documented metabolic depression intertidal bivalves undergo during hypoxia (Storey and Storey, 1990). However, this baseline is higher than the values reported in several earlier studies, where heart rates averaged near zero during emersion (Coleman, 1972; Coleman and Trueman, 1971; Schlieper, 1955). This discrepancy is likely an artifact of our heart rate extraction algorithm, which cannot resolve heart rates below 2 bpm. As evidenced by the drop in extraction success during emersion (Figure 6a), this calculated baseline was likely overestimated. Nevertheless, earlier studies mostly relied on the manual counting from impedance pneumographs (Vereycken and Aldridge, 2023), and a heart rate of 0 bpm may have been subjectively assigned from a seemingly flat signal.

Additionally, the present study recorded a reimmersion overshoot in heart rate lasting about 2 h (Figures 7a and 8a), which is similar in duration to those observed in previous works (Connor and Gracey, 2012; Shick et al., 1986). This overshoot is a well-established mechanism for repaying the oxygen debt incurred during emersion, reflecting the high oxygen demand required to reestablish cellular homeostasis after a period of anaerobiosis (Bayne et al., 1976; Ellington, 1983).

This study also showed that the body temperature of the mussels closely matched the environmental temperature during immersion but became elevated during emersion (Figures 3 and 4). Because this elevation occurred even at night, when the LED strips and heat lamps were inactive, it likely reflects microhabitat temperature variations within the chambers rather than inherent differences in the thermal properties of the robomussels and chamber sensors. Regardless of this elevation, temperature appeared to have limited effects on heart rate during emersion, unlike its strong positive effects during immersion (Figures 7b and 8b). While the high thermal sensitivity of heart rate during immersion is a well-documented phenomenon in *Mytilus* mussels (e.g., Pickens, 1965; Widdows, 1973), the low sensitivity observed during emersion, to our knowledge, has only been reported by Coleman (1973). These findings together suggest that hypoxia-induced metabolic depression effectively overrides the thermal response of heart rate. However, caution is needed before drawing a firm conclusion, given the differential success of heart rate extraction across the temperature range during emersion (Figure 6b). Here, the low extraction success may have led to an overestimation of heart rate at intermediate temperatures, consequently distorting the observed thermal response.

Notably, heart rate extraction success decreased with increasing mussel volume. This is likely due to the greater shell thickness of larger individuals, which reduces the penetration of the infrared light emitted and received by the sensors (Burnett et al., 2013). Additionally, although larger mussels generally have lower heart rates (Pickens, 1965), this negative effect was negligible in the present study. This could be explained by the narrow size range of the mussels selected for the experiment, which limited the ability to detect such allometric scaling. Similarly, the addition of food had a negligible effect on heart rate, which aligns with previous works that found no acute changes associated with feeding (Pickens, 1965; Thompson and Bayne, 1972; Widdows, 1973).

### 4.3 Heart rate after the extreme event

The prolonged emersion event not only led to an overall increase in heart rate but also substantially altered its normal patterns (Figure 8). First, the subtle emersion spike in heart rate disappeared and was replaced by a brief plateau before the eventual decline (Figure 8a). One possible explanation is that the cardiac system was already operating near its upper physiological limit, preventing any further acute increase upon emersion. Second, the reimmersion overshoot was extended, indicating a severe oxygen debt incurred during the event. This finding aligns with Coleman (1972), who reported a positive correlation between the extent of the heart rate overshoot and the duration of the previous emersion. Third, during immersion, the heart rate became more sensitive to temperature changes (Figure 8b), suggesting that environmental heat, which was previously manageable, now posed an additional metabolic burden. This observation exemplifies the interaction between multiple stressors, where exposure to one stressor sensitizes the organisms to another (Sokolova and Lannig, 2008).

Despite these altered patterns, the elevated heart rate steadily decreased in the days following the event, approaching the normal pre-event baseline (Figure 8c). Such returns to baseline have also been reported in mussels after prolonged periods of aquatic hypoxia or hyposalinity (Bakhmet et al., 2005; Klopp, 2025). Together, these findings demonstrate the significant physiological resilience of mussels to environmental stressors.

Ultimately, our work highlights the potential of high-resolution heart rate data in environmental biomonitoring (Vereycken and Aldridge, 2023). Specific derived parameters, such as the general heart rate elevation, reimmersion overshoot duration, heightened temperature sensitivity, and time to return to baseline, could serve as robust metrics to assess the effects of environmental stressors. More broadly, continuous physiological monitoring of intertidal organisms can provide more insights into the consequences of extreme weather events (e.g., heatwaves and floods) in coastal ecosystems (Achterberg et al., 2025).

### 4.4 Conclusions

By continuously monitoring heart rate in the blue mussel *Mytilus edulis* under simulated intertidal conditions, our study confirmed the presence of an often-overlooked emersion spike, as well as the characteristic reimmersion overshoot and differential thermal responses. These heart rate patterns were markedly altered after an unintended prolonged emersion event, resulting in the loss of the emersion spike, an extended reimmersion overshoot, and heightened thermal sensitivity, along with an overall elevation in heart rate. Nevertheless, the mussels exhibited significant resilience, steadily recovering their baseline heart rate in the days following the event. This study demonstrates that continuous physiological monitoring could be useful for assessing the effects of environmental stressors on coastal ecosystems.

## CRediT author statement

**Duy Nghia Pham**: Conceptualization, Methodology, Investigation, Data curation, Software, Formal analysis, Visualization, Writing - original draft.

**Inna M. Sokolova**: Conceptualization, Methodology, Funding acquisition, Resources, Project administration, Supervision, Validation, Writing - review & editing.

## Data availability

The companion R package, rambur v0.5.4, is available on Github at https://github.com/phamdn/rambur/releases#release-v0.5.4. Data and analyses are available on Zenodo at https://doi.org/10.5281/zenodo.10912515.

## Acknowledgments

We thank Shixiu Wang for assisting with animal collection and Holger Pielenz for assembling intertidal chambers. This work was supported by the German Federal Ministry of Research, Technology and Space (BMFTR) through the DAM mission mareXtreme, project ElbeXtreme, grant number 03F0954K.

